# Platelet generation from circulating megakaryocytes is triggered in the lung vasculature

**DOI:** 10.1101/2021.11.01.466743

**Authors:** Xiaojuan Zhao, Dominic Alibhai, Tony G. Walsh, Nathalie Tarassova, Semra Z. Birol, Christopher M. Williams, Chris R. Neil, Elizabeth W. Aitken, Amie K. Waller, Jose Ballester-Beltran, Peter W. Gunning, Edna C. Hardeman, Ejaife O. Agbani, Ingeborg Hers, Cedric Ghevaert, Alastair W. Poole

## Abstract

Platelets, small hemostatic blood cells, are derived from megakaryocytes, although the generation process is not clear. Only small numbers of platelets have been produced in systems outside the body, where bone marrow and lung are proposed as sites of platelet generation. Here we show that perfusion of megakaryocytes *ex vivo* through the mouse lung vasculature generates very large numbers of platelets, up to 3,000 per megakaryocyte. Despite their large size, megakaryocytes were able repeatedly to passage through the lung vasculature, leading to enucleation and fragmentation to generate platelets intravascularly. Using the *ex vivo* lung and a novel *in vitro* microfluidic chamber we determined the contributions of oxygenation, ventilation and endothelial cell health to platelet generation, and showed a critical role for the actin regulator TPM4.

**One-Sentence Summary:** Megakaryocytes form platelets intravascularly in the lung, dependent upon oxygenation, endothelium and megakaryocyte TPM4

---

Platelets are small anucleate blood cells(*1, 2*), ranging from 1 to 3 μm in diameter, with critical roles in haemostasis, thrombosis, inflammation, vascularization, innate immunity and tissue regeneration (*3, 4*). In vitro-derived platelets, as an alternative to native platelets, are attractive for fundamental research because of their rapid genetic tractability, as vectors for drug and genetic component delivery(*5*) and in clinical platelet transfusion. At present, however, their very low production rate, and poor agonist responsiveness, are major obstacles. Platelets are formed by fragmentation from mature polyploid megakaryocytes (MKs), their precursor cells(*6*), although the process of their generation remains incompletely understood(*7, 8*). Bone marrow is proposed to be the main site of platelet production, however indirect evidence since the 1930s(*9, 10*) and recent direct observation(*11*) has shown that MKs can exit the bone marrow and enter the pulmonary vasculature, the first microvascular bed they encounter in the circulation, and that the lung can also be a primary site of platelet biogenesis.

The lung has been proposed as a site of platelet generation by several groups periodically, the most recent evidence presented by Lefrancais et al(*11*). Much of the evidence is indirect, demonstrating the existence of megakaryocytes within lung tissue or vasculature(*9, 11*) and there is now a need to understand the mechanism underlying this platelet generation. For this reason, we established an *ex vivo* mouse heart-lung model (Fig. 1A) through which we were able to perfuse murine MKs. Remarkably, we could show for the first time that MKs, despite their size, can pass multiple times through the lung vasculature, and that this leads to the generation of physiological levels of functional platelets (approximately 1,000~4,000 per megakaryocyte(*7, 12*)). The study investigates the mechanism of platelet generation in the lung vasculature and shows that air ventilation and healthy pulmonary ECs play critical roles for platelet generation, suggesting the lung vasculature as a primary site of platelet biogenesis. We show for the first time that MKs undergo enucleation upon repeated passage through pulmonary vasculature before cytoplasm fragmentation to generate platelets. This final process of fragmentation of enucleated MKs depends upon expression of the actin regulator TPM4, suggesting an active cytoskeleton-dependent mechanism in the lung vasculature.

**Fig. 1:**
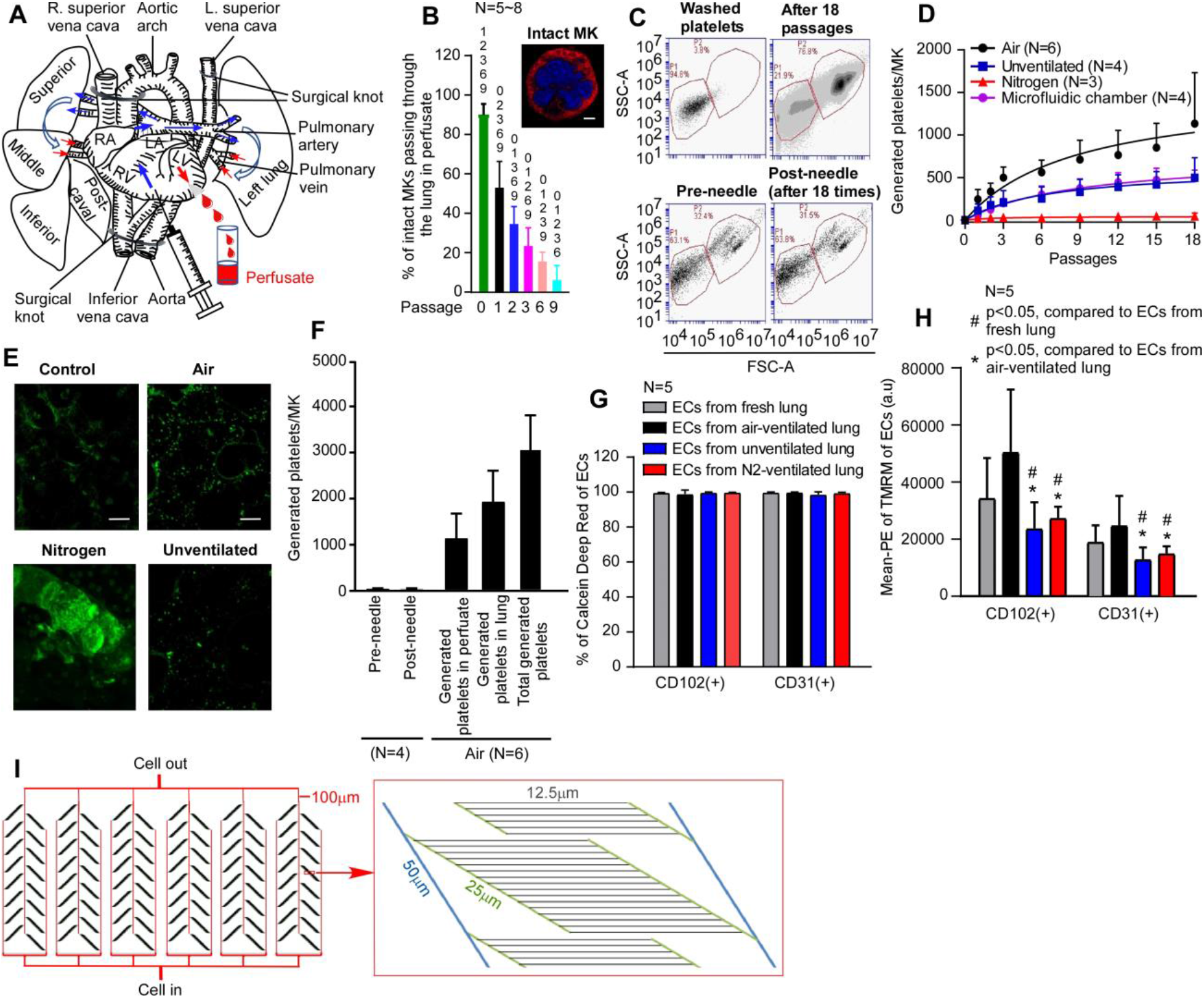
Mouse platelets are generated in physiological quantities from megakaryocytes passaged multiple times through mouse pulmonary vasculature *ex vivo*. **(A)** Diagram illustrating the approach to generating mouse platelets from megakaryocytes (MKs) by infusion through the heart-lung preparation. Inferior and superior venae cava, plus the aortic arch, were ligated (indicated as surgical knot), and the MK suspension was pumped into the pulmonary vasculature through the right ventricle (RV). The perfusate was collected via the left ventricle (LV), as indicated. **(B)** Approximately 3×10^4^ mouse MKs, labelled with PE-conjugated anti-CD41 antibody, were passaged through the pulmonary vasculature *ex vivo*. Intact MKs (inset: showing a circular shape and central nucleus, scale bar: 10 μm) were imaged from samples of perfusates after passage numbers 1, 2, 3, 6 & 9 through murine lung vasculature ex vivo. Quantification was from at least 250 fields of view, counting at least 230 cells in total and displayed as a percentage of total number of cells. Numbers above each column are passage number against which the column show a significant difference, as determined by one-way ANOVA test. P-value less than 0.05 was considered statistically significant. Data are from at least 5 independent experiments. **(C)** Gating strategy for quantification of generated platelets. Washed mouse platelets were used to set gates for generated platelets (P1) at the FSC/SSC density plot by size and granularity. MKs were passaged 18 times through the pulmonary vasculature *ex vivo*. The number of generated platelets in the perfusate collected after the 18^th^ passage was determined by the number of CD41(+) events in gate P1. MK suspensions, either non-passaged (pre-needle) or passaged through a 21G needle 18 times (post-needle), were evaluated to exclude whether multiple needle passages would be sufficient for MKs to generate platelets. **(D)** The number of generated platelets per MK present in the perfusates from different passage numbers in lungs either ventilated with air, 100% nitrogen, or without ventilation (as indicated), or from microfluidic chamber (shown diagrammatically in **I**) were measured by FACS. Data are mean and S.E.M. (n as indicated). **(E)** Stained MKs were passaged through pulmonary vasculature *ex vivo* 18 times, and lung tissue visualized at 20 stacked focal planes by two-photon microscopy. During the cell passage lungs were either ventilated with air or 100% nitrogen, or were not ventilated, as indicated. Mouse lung without MKs passed through served as control. Images shown are representative of at least 4 independent experiments. scale bar: 20 μm. **(F)** Numbers of platelets generated per MK in perfusate and remaining in mouse lung (as calculated from two-photon imaging per volume, extrapolated to the total lung volume) under air ventilation, were calculated and displayed as means +/−S.E.M (n as indicated). As a control, MKs passed through 21G needles 18 times generated no platelets, as indicated. (**G-H**) The viability (**G**) and mitochondrial membrane potential (**H**) of pulmonary endothelial cells (ECs) were measured by FACS. Pulmonary ECs were isolated from perfused lungs under air- or pure-N_2_-ventilation or unventilated for approximately 2 hours and marked with either anti-CD31/PECAM-1 or anti-CD102/ICAM-2 antibodies. Pulmonary ECs from fresh lung served as control. Comparison was by unpaired student *t*-test. P-value less than 0.05 was considered statistically significant. Data are from 5 independent experiments. (**G**) The viabilities of pulmonary ECs were determined by Calcein Deep Red retention and displayed as means of % +/−S.E.M. (**H**) The mitochondrial membrane potential was determined by accumulation of tetramethyl rhodamine methyl ester (TMRM) in active mitochondria and displayed as mean intensity +/−S.E.M. (**I**) Design of microfluidic chamber. A set of channels mimicking a physiological vascular system was constructed using standard PDMS approach. The design shows a branching structure such that as branching progressing, the channel diameter decreases by half. All channels are rectangular in cross-section and 10 μm deep, with the largest channels being 100 μm across, decreasing to the smallest channels which are 12.5 μm across. From each larger channel, 16 smaller channels branch off, allowing for maintenance of flow resistance due to the r^4^ power relationship between resistance and channel diameter. Fluid then flows from larger diameter channels to smaller diameter channels, and in reverse on the way out of the system. Cells are passed through the system, repeatedly. The system is scaled up, through multiplexing in parallel, to allow greater cell volumes to be used, as shown.

## Platelet generation in the *ex vivo* lung vasculature

The *ex vivo* mouse heart-lung model (Fig. 1A) is based upon isolating the heart and lungs as a single unit, ligating the venae cavae and the aortic arch and perfusing CD41-PE or FITC-labelled MKs through the pulmonary circulation from the right ventricle, collecting the perfusate from the left ventricle. This allows quantitation and imaging of cells passaged through the pulmonary vasculature. In the first instance, lungs were artificially ventilated with air. We were expecting that the vast majority of MKs would be trapped within the lung vasculature due to their large size (around 50-100 μm diameter(*13*)), but unexpectedly more than 50% of the MKs emerged in the perfusate after the first passage (Fig. 1B). The perfusate could be re-injected through the lungs and although upon multiple passages the number of intact MKs (showing a circular shape and central nucleus) continued to decrease, it was evident that MKs were able to pass through the lung vasculature multiple times. It was also apparent that CD41-labelled particles also appeared in the perfusate that were similar in size and granularity to mouse platelets. We term these “generated” platelets (CD41-positive events in gate P1 in Fig. 1C). The numbers of generated platelets per MK gradually increased with increasing passages, up to around 1134.7±597.2 platelets/MK after 18 passages (Fig. 1D and F). Two-photon microscopy of fixed lung sections after 18 passages showed that some generated platelets were retained in the lung microvasculature (Fig. 1E and Movie S1-3). These were quantified in a defined lung volume and the total lung volume measured using a fluid displacement method (Fig. S1B) to estimate numbers of retained platelets. We calculated this to be 1950±682 per MK injected (Fig. 1F). Adding this to the number contained in the perfusate (Fig. 1D), we estimate that after 18 passages through the lung approximately 3084.7±1279.2 platelets were generated per MK (Fig. 1F), in keeping with previous estimates that each MK *in vivo* produces approximately 1000-4000 platelets(*7, 12*). Platelets were not generated simply as a consequence of passage through small-bore needles, since we showed that repeated passage of MKs through 21G needles 18 times is not sufficient to generate platelets (Fig. 1C and Fig. 1F). Altogether, we could generate physiological level platelets after multiple recirculation of MKs through the lung microvasculature. We also conclude that MKs substantially, rapidly and reversibly, deform their shape to passage through the capillary network of the lung.

## Essential role for air ventilation and healthy pulmonary endothelial cells in platelet generation in mouse lung

The *ex vivo* mouse heart-lung model (Fig. 1A) can be a useful tool to allow artificial ventilation with either ambient air or with pure nitrogen (N_2_), or no ventilation, to assess the roles of physical ventilation and gaseous oxygen in regulating MK biology and thrombogenesis.

We first explored whether air ventilation is essential for platelet generation in our model. Strikingly, when lungs are ventilated with pure N_2_ to completely de-oxygenate the heart-lung preparation, the number of generated platelets in the perfusate was almost ablated, reduced to just 43 ± 20 platelets/MK (Fig. 1D). Two-photon imaging of nitrogen-ventilated lungs showed mature MKs were trapped in the lung vasculature (Fig. 1E and Movie S4), a feature not observed under air ventilation. Pulmonary endothelial cells (ECs), which play key roles in gas exchange in the lung(*14*), interact closely with MKs as they passage through the vasculature, and we therefore compared their viabilities (determined by the Calcein Deep Red retention assay) and mitochondria membrane potential (an indicator of healthy cells, determined by accumulation of tetramethyl rhodamine methyl ester (TMRM) in active mitochondria) after 18 passages under air- or N2-ventilation for approximately 2 hours. Surprisingly, ECs from preparations ventilated under N2 were fully viable, and comparable with those ventilated under air (Fig. 1G). However, the mean intensity of TMRM of ECs from N2-ventilated lungs was approximately half that of air-ventilated lung or fresh lung (Fig. 1H). These data suggest that air ventilation and fully healthy ECs are essential for efficient platelet generation in the heart-lung preparation, and the capability of MKs to pass through the lung vasculature is lost in heart-lung preparations that were N2-ventilated. We then investigated whether healthy pulmonary ECs are required for the passage of MKs through the heart-lung preparation and the importance of ventilation for platelet generation. After flushing the lung under air ventilation, we then stopped ventilation while perfusing MKs through the preparation 18 times. In the absence of ventilation, the number of platelets generated per MK in the perfusate still gradually increased with increasing passage number, but the numbers generated were substantially lower than in the air-ventilated condition (498.4± 235.7 platelets/MK, Fig. 1D). Importantly, the mean intensity of endothelial cell TMRM was approximately halved by lack of ventilation relative to air-ventilated controls, equivalent to the N2-ventilated lung (Fig. 1H). In contrast to the N2-ventilated lung however, two-photon imaging of unventilated lungs showed no MKs trapped in the lung vasculature (Fig. 1E). The data suggest therefore that passage of MKs through the pulmonary vasculature may not require a fully healthy endothelium, but clearly oxygenation plays a critical role at some level, since MKs cannot passage through the vasculature in N2-ventilated lungs. This effect may reflect a direct requirement of MKs for oxygenation, but the data also suggest that the passage of MKs through structural arrangement of the lung microcirculation is also a major contributor to platelet generation.

To verify whether this structural arrangement could mediate to platelet generation, we designed a polydimethylsiloxane (PDMS)-based (gas permeable) microfluidic chamber with channel arrangement mimicking tissue microcirculation. The channels were of uniform depth of 10 μm throughout, where the entry and exit channels had a width of 100 μm and where branches emerged halving the channel width each time, to a minimal width of 12.5 μm, as per the diagram shown in Fig. 1I and Fig. S1C. This channel arrangement allowed us to flow through cells and determine platelet generation in the perfusate after repeated passage. Fig. 1D shows that the numbers of generated platelets per MK gradually increased with increasing passages, similar to the numbers generated in the unventilated lung, with 492.3±95.1 platelets/MK after 18 passages (Fig. 1D). Altogether, these data suggested that (1) air-ventilation and healthy ECs are required for MKs to generate physiological levels of platelets in the heart-lung preparation; (2) the structural arrangement of the pulmonary microcirculation plays a role in platelet generation; (3) lack of ventilation or N2-ventilation for 2 hours caused partial loss of the mitochondrial membrane potential of pulmonary ECs; (4) the exposure of pure N_2_ for 2 hours impaired the mobility of megakaryocytes.

## Generated platelets are morphologically and functionally normal

We determined that generated platelets were *bona fide* live platelets rather than cellular fragments, using the vital dye calcein-AM (Fig. 2A) and showing mitochondrial function comparable to normal mouse platelets using TMRM (Fig. 2B-C). Generated platelets were anuclear, showing no staining with the DNA dye Draq5 (Fig. 2A). The proportion of cells expressing CD61 and CD42b, and the mean fluorescence intensity of those markers, were comparable between generated and control platelets (Fig. 2D). However, the mean fluorescence intensity of three collagen receptors GPVI, CD49b(*15*) and CD42d(*16*) was higher in generated platelets than controls (Fig. 2D), while the proportion of cells expressing these receptors was lower (Fig. S1D). Platelets display an almost uniquely characteristic sub-plasma membrane microtubular ring, running circumferentially in resting platelets(*17, 18*). Our generated platelets, immunolabelled for α-tubulin, display this characteristic ring structure (Fig. 2E), and the cells are larger than controls (3.5 ± 1.2 μm vs 1.9 ± 0.3 μm, Fig. 2F). Younger platelets have been shown to be larger than the mean platelet diameter(*19, 20*), and our generated platelets are therefore consistent with this observation. Finally, we visualized the ultrastructures of generated platelets by transmission electron microscopy (TEM, Fig. 2G), after depletion of host platelets using anti-GPIbα antibodies. Generated platelets displayed a discoid shape with classical characteristics including α-granules, σ-granules or dense bodies, mitochondria, open canalicular system, and microtubule coils.

**Fig. 2:**
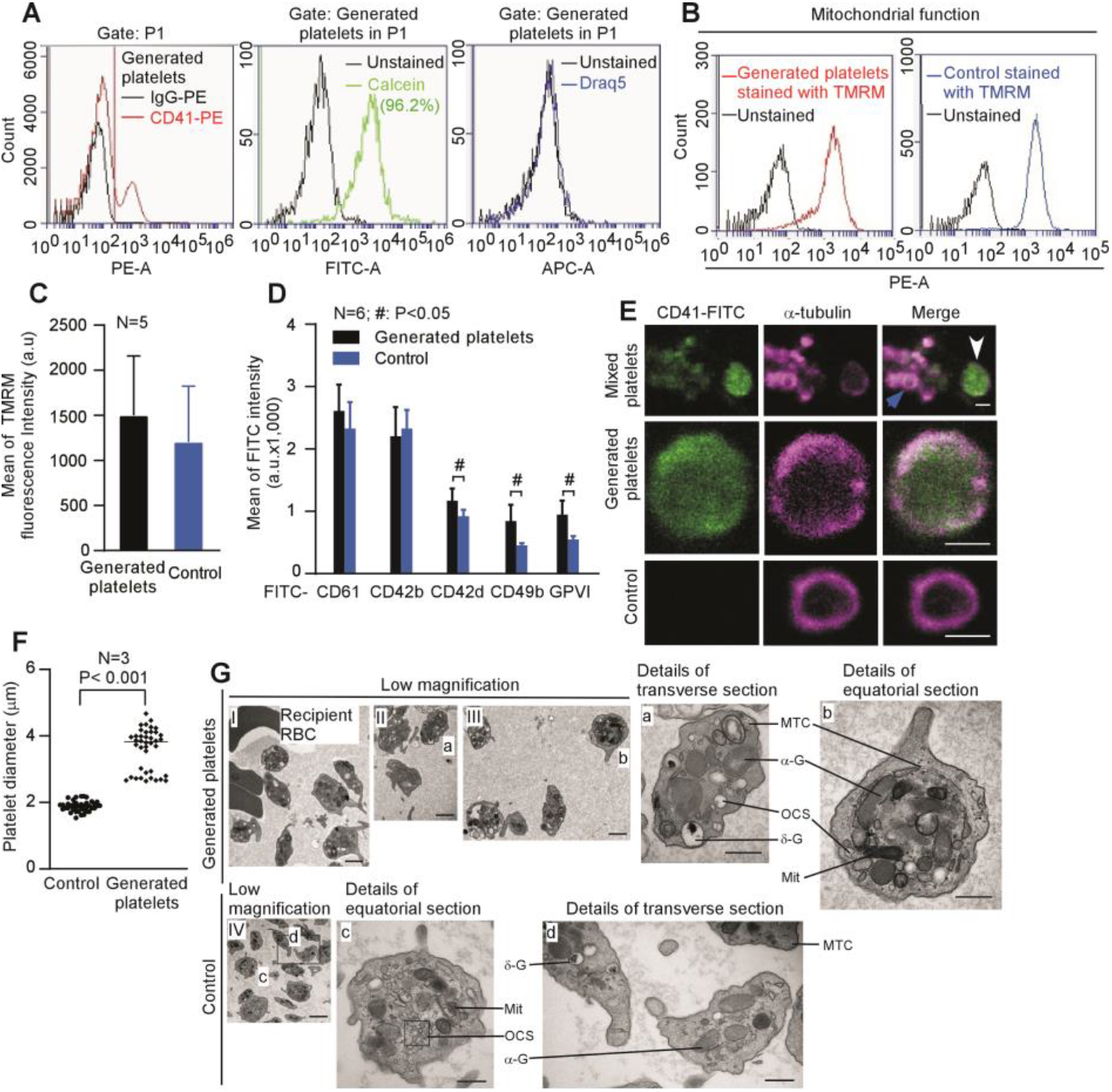
Generated platelets demonstrate typical physical features comparable to control platelets. **(A)** Platelet populations in P1 (red) were separated into two subtypes: (i) events with similar mean fluorescence to those derived from control IgG-PE treated MKs (black), which are comprised mainly of control (host) platelets and (ii) events with higher mean fluorescence compared to those derived from control IgG-PE-treated MKs, which are the generated platelets. The viability of the generated platelets, and whether they contain DNA, were checked by Calcein AM and DRAQ5 dyes, respectively, and presented as line graphs. **(B)** Mitochondrial membrane potential was monitored by FACS in generated and control platelets after loading with TMRM dye. **(C)** TMRM signals from (B) were quantified and displayed as mean +/−S.E.M (n=5). **(D)** Platelets were defined by staining with anti-CD41 PE conjugated antibody. Surface glycoproteins of platelets were measured by FACS on generated platelets and control platelets after staining with different FITC-conjugated antibodies as indicated. Data are presented as mean +/−S.E.M (n=6) of mean FITC fluorescence intensities. **(E)** Perfusates, containing both generated platelets (stained green with CD41-FITC, white arrow) and host platelets, were stained for α-tubulin (stained magenta, blue arrow), and confocal images shown as a mixed population in the top panels. More detailed images of a-tubulin rings are shown in the magnified images in the middle panel (generated platelets) and bottom panel (control platelets). Images are representative of 3 independent experiments. Scale bars: 2 μm. **(F)** The diameter of platelets (40 platelets from 3 independent experiments) from (E) was measured using Fiji (ImageJ-Win64). Comparison was by unpaired student *t*-test. P-value less than 0.05 was considered statistically significant. **(G)** Ultrastructures of generated platelets and control platelets visualized by transmission electron microscopy (TEM). Host platelets were first depleted by intraperitoneal administration of anti-GPIbα antibody R300 (2 μg/g bodyweight) prior to MKs infusion through the heart-lung preparation. Low magnification images are shown for generated platelets in I-III, and control platelets in IV. Detailed images, taken from the low magnification images, are shown in (a)-(d), as indicated. Subcellular structures are shown and annotated as abbreviations: α-G, α-granules; σ-G, σ-granules or dense bodies; Mit, mitochondria; OCS, open canalicular system; MTC, microtubule coils; RBC, red blood cells. Scale bars: 2 μm in I-IV, 500 nm in a-d. Images shown are representative of 5 independent experiments.

We then demonstrated that the functionality of generated platelets is also similar to that of mouse platelets (controls). Both generated and control platelets showed equivalent responses to agonists (CRP and thrombin) in terms of integrin α_IIb_β_3_ activation and degranulation (P-selectin expression, Fig. 3 A-B). Thrombus formation *in vitro* was also assessed, determining how generated platelets mixed into whole blood interact with a collagen-coated surface under flow. Generated platelets (stained with both DiOC6 and CellTracker™ Red CMTPX dye, white) occupied all levels of the thrombus whilst control platelets (stained with CellTracker™ Red CMTPX dye alone, magenta) were mainly situated on top of the thrombus, suggesting generated platelets showed a higher responsiveness to collagen, or were primary reactors to it (Fig. 3C-E and Movie S5). This may suggest that generated platelets are early interactors with collagen, displaying the greater adhesive functionality of younger platelets(*19, 20*), possibly due to higher levels of collagen adhesive receptors (GPVI and CD49b(*15*), and CD42d(*16*)) (Fig. 2D).

**Fig. 3:**
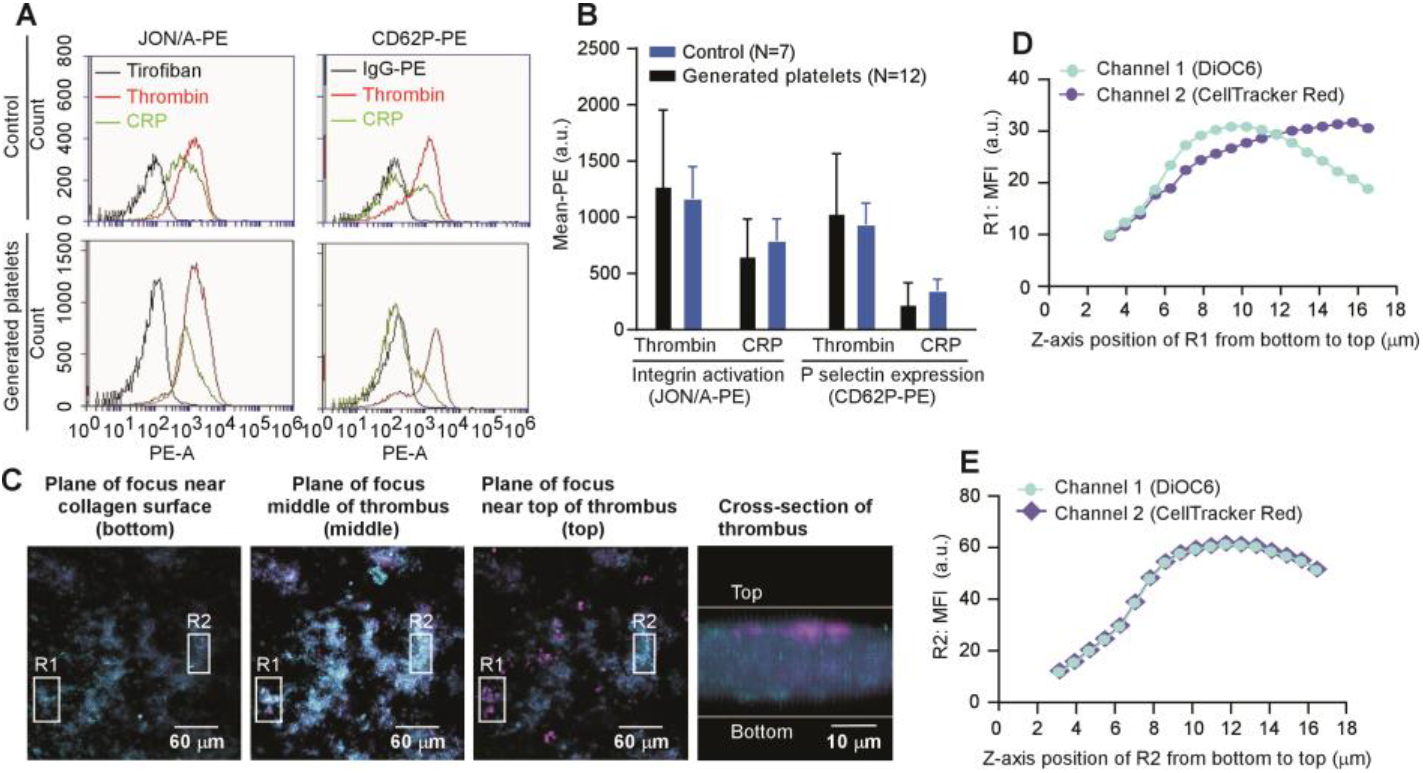
Generated platelets are functionality comparable to control platelets. Mouse MKs, labelled with FITC-conjugated anti-CD41 antibody, were passaged repeatedly through the pulmonary vasculature *ex vivo*. Lungs were ventilated with air throughout **(A)** Representative FACS histograms of integrin αII_b_β_3_ activation (JON/A binding) and P-selectin expression induced by 2 U/ml thrombin (Magenta) or 5 μg/ml CRP (green) in washed generated platelets under air ventilation in comparison to washed control platelets. Tirofiban (80 ng/ml) was used as control for non-specific binding in measurements of integrin activation. **(B)** Data from (A) were expressed as mean fluorescence intensity (a.u.). Platelets generated under air ventilation showed similar integrin αII_b_β_3_ activation and P-selectin expression in response to thrombin and CRP compared to control platelets. Data are mean and S.E.M. (n as indicated); comparison by Student’s unpaired, two-tailed *t*-test. **(C)** Images of 3 planes of a representative platelet-rich thrombus, formed by flowing of blood at arterial shear (1000s^−1^) over a collagen-coated glass surface. Cross-sectional image is shown on the far right. Mouse MKs were stained with DiOC6 and passed through mouse lung 18 times. The perfusate was then mixed with mouse blood from the host, from which platelet rich plasma (PRP) had been removed. Blood was then loaded with CellTracker™ Red CMTPX dye. Generated platelets (stained with both DiOC6 and CellTracker™ Red CMTPX dye, white) occupied all levels of the thrombus while host platelets (stained with CellTracker™ Red CMTPX dye alone, magenta) were mainly situated on top of thrombus. Images are representative of 5 independent experiments. Scale bars as indicated. **(D-E)** MFI (mean fluorescence intensity) profiles of R1 and R2 (Region 1 and RegioN2 from Fig. 3C) along the z-axis. **(D)** MFI profile of R1 along the z-axis. In R1, generated platelets occupied the lower part of the thrombus up to ~12 μm (both green and meganta signals increased simultaneously), whilst beyond this point host platelets were predominant (meganta signals were stronger than green beyond 12 μm). **(E)** MFI profile of R2 along the z-axis. In R2, this part of thrombus was composed only of generated platelets as both green and meganta signals changed simultaneously along z-axis.

## Multiple passage of megakaryocytes through lung: enucleation and platelet generation

We wanted to explore the details of the release of platelets from MKs upon repeated passage through the pulmonary vasculature. The cells in the perfusate were imaged after collected over a defined numbers of passages (3, 6, 9, 12 and 18), and strikingly, upon repeated passages, MKs gradually move their nuclei to the periphery and subsequently enucleate, generating both naked nuclei and enucleated MKs. Although small numbers of enucleated round MKs were found, we saw the gradual accumulation of larger anuclear objects. Figure 1B, 4 and Movie S6-8 show the steps involved in the process, with images shown in Fig. 4A, quantified in Fig. 1B and 4B and summarized in Fig. 4J. As shown in Fig. 1B, the percentage of intact MKs decreased from 54% after 1 passage (P1) to 6.7% after 9 passages (P9), while in Fig. 4B the percentage of large naked nuclei (>20 μm diameter) increased from 5% after P1 to 22% after P9. At the same time, large anuclear objects (>10 μm) also increased, as a proportion of total MKs and derivatives, from 13% after P1 to 45% after P9. These data suggested that the large polyploid nucleus moves from a central position to the periphery of the cell, in a process of polarization. The nucleus is then extruded from the cell upon further passages through the lung vasculature, until by approximately 9 passages there are very few nucleated MKs left. After 12 passages, large naked nuclei (>20 μm) also became rare, being replaced by irregular small sub-nuclei. These sub-nuclei appeared connected to each other, probably by membranous structures which were too fine for visualization by light microscopy (Fig. 4C and Movie S9). Sub-nuclei were characterized by depth (Fig. 4D) and aspect ratio (width: height, Fig. 4E), all of which decreased substantially with increasing passage number (P3 to P18 parameters: depth 8.9 μm to 5.3 μm, aspect ratio 2 to nearly 1, major axis 14 μm to 6.3 μm, minor axis 8 μm to 5.7 μm). The extruded naked nuclei therefore undergo a process of division into multiple component sub-nuclei, which proceed to condense into compact sub-nuclear units with a greater circularity. The anucleate MK proceeds to fragment into platelets after multiple passages, to reach plateau numbers by 15-18 passages (see Fig. 1D). This is the first time these enucleating behaviors have been observed for MKs when releasing platelets.

**Fig. 4:**
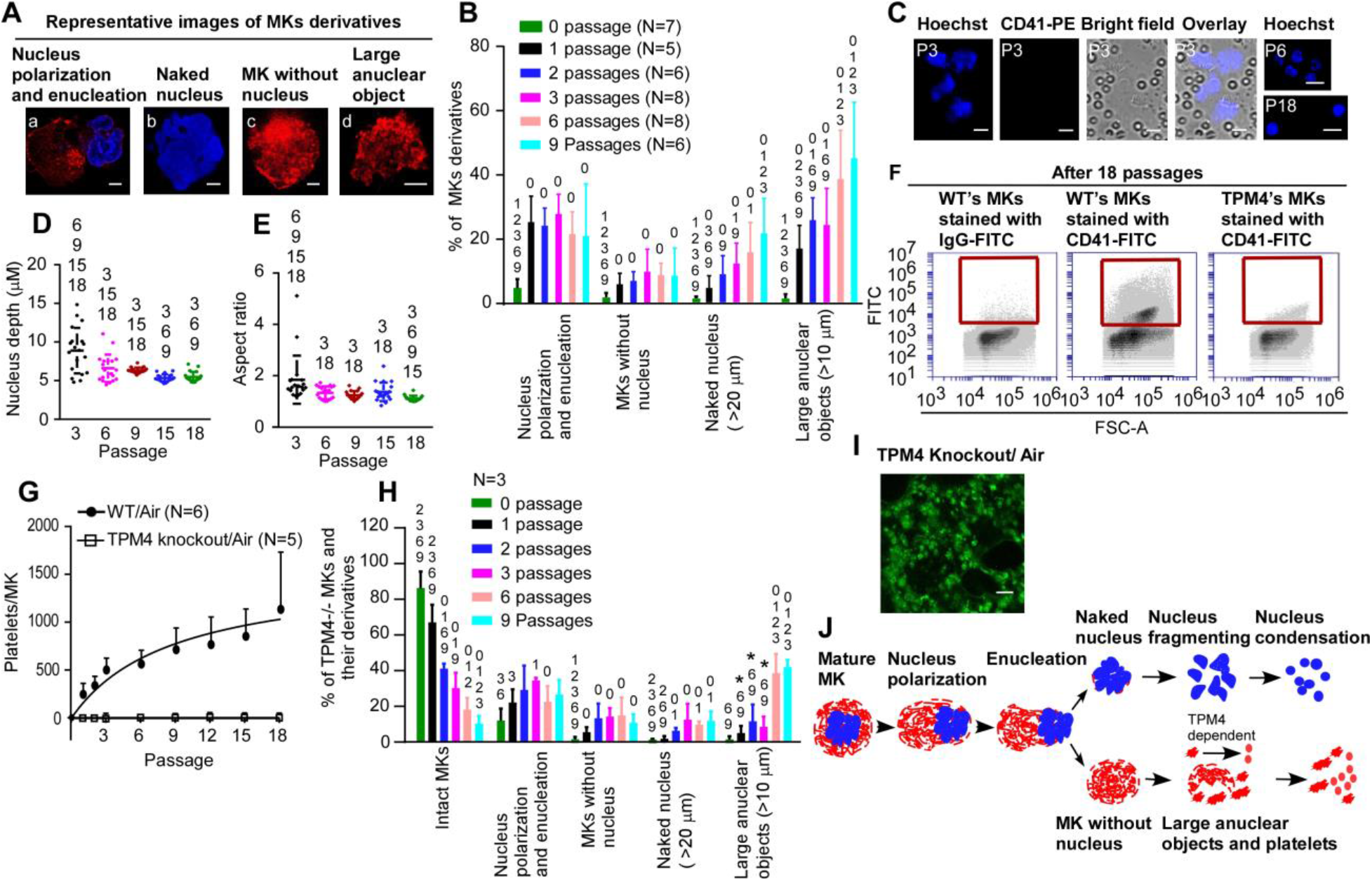
Megakaryocytes show nuclear marginalization and enucleation prior to TPM4-dependent fragmentation. **(A-E)** MKs from C57BL/6 mice, labelled with PE-conjugated anti-CD41 antibody, **(F-I)** MKs from *Tpm4*^−/−^ mice, labelled with FITC-conjugated anti-CD41 antibody. MKs were passaged repeatedly through the pulmonary vasculature of a C57BL/6 mouse *ex vivo*. Lungs were ventilated with air throughout. **(A)** Representative images of MK derivatives during the process of platelet generation: (a) <u>nuclear polarization and enucleation</u>, where the nucleus is marginalized, of irregular shape or in the process of ejection from the cell; (b) <u>naked nuclei</u>, where the ejected nucleus is larger than 20 μm in diameter and free from the parent cell and may be partially encased in a thin cytoplasmic layer; (c) <u>MKs without nuclei</u>, where the MKs have an approximately circular shape; and (d) <u>large anuclear objects</u>, where ghost cells are of irregular shape and larger than 10 μm in their longer axis. **(B) & (H)** Cells were imaged from samples of perfusates after passage numbers 1, 2, 3, 6 & 9 through murine lung vasculature *ex vivo*. Cells were morphologically classified as 5 subgroups: intact Mks (Fig.1B) and (a)-(d) above and quantified as a percentage of total number of cells. Quantification was from at least 250 **(B)** or 150 **(H)** fields of view, counting at least 230 **(B)** or 170 **(H)** cells in total for each of the subgroups. Numbers above each column are passage numbers against which the column shows a significant difference (p<0.05), as determined by one-way ANOVA test. Asterisk above columns in (H) represents significant difference to corresponding wild-type column in **(B)**. Data are from at least 5 **(B)** or 3 **(H)** independent experiments. **(C-E)** Nuclear lobes of MKs fragment into small condensed sub-nuclei. **(C)** Representative images, from n=3, of naked nuclei generated after multiple passages (as indicated) of mouse MKs through pulmonary vasculature. Scale bars: 5 μm. **(D-E)** The depth **(D)** and aspect ratio **(E)** of sub-nuclei decreased substantially with increasing passage number. For depth **(D)**, the decrease was from ~8.9 μm after P3 to 5.3 μm after 18 passages (P18). For aspect ratio **(E)**, the decrease was from ~2 after P3 to nearly 1 after P18. Each symbol represents one sub-nucleus. Numbers above each column are passage numbers against which the column shows a significant difference, as determined by one-way ANOVA. P<0.05 was considered significant. Data are from 3 independent experiments. **(F-G)** Numbers of generated platelets per *Tpm4*^−/−^ MKs, or control WT MKs, in perfusates after different passage numbers through WT lung, were quantified by FACS. **(F)** Representative FACS dot plot images for generated platelets (CD41-FITC positive events within the red square). **(G)** *Tpm4*^−/−^ platelets were consistently undetectable after up to 18 passages, in the perfusate. Data shown are platelets generated per MK from either *Tpm4*^−/−^ MKs or control WT MKs and displayed as mean +/−S.E.M. (n as indicated). **(H)** see above, with **(B)**. **(I)** Abundant fluorescent objects, 10-12 μm diameter, were visible in sections of mouse lung after 18 passages of stained *Tpm4*^−/−^ MKs, as shown in extended focus stacks of 20 continuous two-photon planes of lung. Images shown are representative of 3 independent experiments. Scale bar: 20 μm. **(J)** Schematic diagram of the steps in platelet generation from mature MK. These include nuclear polarization, enucleation, cytoplasmic fragmentation into platelets and nuclear condensation and fragmentation.

## Tropomyosin 4 required for platelet generation in the lung vasculature

The release of platelets is understood to require cytoskeletal re-organisation involving the actin cytoskeleton. Tropomyosins form co-polymers with actin filaments and regulate filament function in an isoform-specific manner(*21*). Tropomyosin TPM4 has been shown to have a role in platelet formation^14^. In our *ex vivo* lung system, *Tpm4^−/−^* MKs generate no platelets (Fig. 4F-G), in contrast to the *in vivo* setting where platelet counts in *Tpm4*^−/−^mice drop by approximately 50% compared to wild-type(*22*). This suggests that the remaining 50% platelet generation may take place in the bone marrow and may not fully require TPM4. This is consistent with the findings of LeFrancais *et al*.(*11*), who have shown that approximately 50% of circulating platelets are generated in the mouse lung. We then wanted to explore further the steps in platelet generation requiring TPM4. Compared to wild type MKs, during the first 3 passages, fewer *Tpm4*^−/−^ MKs underwent transformation to large anuclear objects. However, this represented only a delay in these events, since by 6-9 passages the numbers of enucleated cells were comparable with wild-type (Fig. 4H). Two-photon microscopy of fixed lung sections after 18 passages showed abundant anuclear fluorescent objects, sized 10 μm (Fig. 4I and Movie S10). These data suggest a small and non-essential role for TPM4 in regulating enucleation, but a critical role in regulating both fragmentation of MKs into platelets with normal sizes (summarized in Fig. 4J).

## Discussion

This study is the first to demonstrate platelet generation from cultured MKs outside the body with an output similar to *in vivo* estimated MK capabilities. Several unexpected findings were observed during the study. First, approximately 50% of MKs pass through the lung microvasculature. This suggests a substantial ability to reversibly deform as they passage through the lung microvasculature. Pulmonary capillaries have average internal diameters of 5-8 μm(*23*) which decrease with lung inflation(*24*). This challenges an assumption that MKs may be trapped in the lung microvasculature, despite the mismatch in sizes of cell and capillary diameter(*25*). However, longer term exposure to pure N_2_, to completely de-oxygenate the lung, over two hours, effectively ablated platelet formation, and caused MKs to lodge in the lung vasculature which seemed mainly due to an impaired MK motility, given that MKs still could pass through the unventilated lung vasculature or microfluidic chambers. MK numbers are increased in the lungs of patients with acute respiratory distress syndrome caused by COVID-19, for example, and pulmonary MK levels may represent a histologic variable of interest in this disease(*26*). Although some of these MKs may be resident cells in the stroma of the lung(*27*), it is possible that they are MKs trapped in lung vasculature, just as we see for lungs ventilated with N_2_.

Second, repeated passage of MKs through the pulmonary vasculature induces a reproducible sequence of events leading to platelet generation. Our data suggest that O2 tension and maintaining healthy pulmonary ECs, as well as repeated physical contraction of the lung are important to support platelet generation. This might suggest an essential and dynamic conversation between the pulmonary microvasculature environment and MKs, we term ‘education’, to induce reversible deformation of MKs, stimulate motility through the microvasculature and promote platelet release. This education makes the lung a primary and unique site for platelet biogenesis. However, since MKs pass through the pulmonary vasculature, it is possible that an important educational signal is induced simply by passage through the small channels of the vasculature, and therefore that some platelet generation may take place as they passage through other organs of the body. This is supported by the observation that we can generate substantial numbers of platelets by passage of MKs through the microfluidic chamber that mimics microvasculature. These pieces of evidence though suggest that there is synergy between all the unique features of the lung vasculature that makes this organ likely to be the most important site of vascular production of platelets, since the production of platelets in the ex vivo lung is greater than in the microfluidic chamber, and suggests important roles for oxygenation, mechanical ventilation and signals from the pulmonary vascular endothelium.

Besides MKs themselves, our data suggest that there are four factors which contribute to platelet generation, including oxygenation, ventilation, healthy pulmonary ECs and the microvascular structure. This may partially explain differences in platelet count reported in patients with lung disease(*28, 29*), or people living at the different altitudes(*30, 31*). These differences may be because the process of platelet generation, described here, is complex with inputs from the four factors described and possible redundancy or partial redundancy between these four factors. For example, there may be compensation for low oxygenation by an increase in respiratory rate(*28*). Third, MKs enucleate intravascularly, as part of the process leading to platelet generation. Few cells are known to enucleate, but importantly these include erythrocytes. This may be important because MKs and erythrocytes are developmentally closely related and share a common precursor, the MK/erythroid progenitor (MEP)(*32, 33*). It is possible therefore that there may be shared machinery important for enucleation in both related cell lineages. Our observation could be consistent with the phenomena that naked nuclei of denuded megakaryocytes were found in the bone marrow and the lung of patients with severe Covid-19, although in healthy individuals these are substantially less common due to rapid removal by the mononuclear phagocyte system(*34*). Fourth, another striking observation was that in the *ex vivo* lung preparation, *Tpm4*^−/−^ murine MKs generated no platelets. Mechanistically we showed this to be by abolishing the fragmentation of MKs into normal sized platelets and their subsequent release from the lung vasculature. In addition, this demonstrated not only a critical role for TPM4 in the MK education process in the lung, but also suggested that platelet generation by lung education may be by a different cellular and molecular mechanism to platelet generation in the bone marrow. These observations are consistent with Lefrancais *et al(11)* who had proposed that approximately 50% of platelet generation takes place in the lung vasculature. The findings suggest that the two systems of platelet generation, in lung and bone marrow, may each operate, and that at least in the mouse each system contributes approximately half of the circulating platelets. Studies in genetically modified mice and correlates with genetic-based causes of thrombocytopenias in humans show that both microtubules(*6*) and actin(*8, 35, 36*) play important roles in platelet generation from MKs. TPM4 has a prominent role in actin regulation, therefore our data suggest that the lung vasculature educates MKs to generate platelets by an actin-based mechanism.

This work identifies a mechanism of platelet generation, by repeated passage of megakaryocytes through lung vasculature under air ventilation, involving enucleation and TPM4-dependent fragmentation of cells. The findings add to our understanding of why the lung is a primary site for platelet generation. Importantly though, this may help to set up new approaches to large scale generation of human platelets, valuable for establishing novel ways to understand the molecular control of platelet generation and function, and for developing a variety of cell-mediated therapies and supply of donor cells for transfusion.

## Acknowledgments

We gratefully acknowledge the Wolfson Bioimaging Facility for their support and assistance in this work. We are also grateful to Professor Jack Mellor, University of Bristol, for the use of the tissue slicer machine.

## Author contributions

Conceptualization: XZ, CG, AWP

Methodology: XZ, DA, TGW, NT, SZB, CRN, EWA, AKW, J B-B

Investigation: XZ, AWP

Visualization: XZ, DA, CRN

Funding acquisition: EOA, CG, AWP

Project administration: AWP

Supervision: XZ, AWP

Writing – original draft: XZ, DA, CRN, AWP

Writing – review & editing: XZ, DA, CMW, PWG, ECH, EOA, IH, CG, AWP

## Data and materials availability

The datasets generated during and/or analyzed during the current study are available from the corresponding authors on reasonable request.

## Supplementary Materials

Materials and Methods

Fig. S1

Movies S1 to S10

## This file includes

Materials and Methods

Fig. S1

Captions for Movies S1 to S10

## Materials and Methods

### Materials

Cell culture reagents including IMDM-Glutamax and penicillin-streptomycin were from Invitrogen (Paisley, UK). Recombinant murine stem cell factor (rSCF) and recombinant murine thrombopoietin (rTPO) were purchased from PeproTech EC (London, UK). PE or FITC-conjugated anti-CD41 and isotype-nonspecific IgG were from BD Biosciences (Berkshire, UK). FITC-conjugated anti-mouse CD31, CD102 antibodies and isotype-nonspecific IgG were from Biolegend (San Diego, USA). SYLGARD™ 184 Silicone Elastomer Kit was from Dow Corning (Michigan, USA). Anti-mouse GPIbα antibody R300, FITC-conjugated anti-mouse CD61, CD42b, CD42d, CD49b, Glycoprotein VI (GPVI) antibodies and isotype-nonspecific IgG, PE-conjugated anti-CD62P antibodies, JON/A anti-integrin α_IIb_β_3_ and isotype-nonspecific IgG were from Emfret Analytics (Würzburg, Germany). Hoechst 33324, DRAQ5, Calcein AM, Calcein Deep Red, tetramethylrhodamine (TMRM) and CellTracker™ Red CMTPX dye were from Molecular Probes™ (Loughborough, UK). DiOC_6_ iodide was supplied by Enzo Life Sciences (Exeter, UK). Ibidi slides were from Thistle Scientific (Glasgow, UK). Horm fibrillary collagen was from Takeda Pharma (Linz, Austria). Cross-linked collagen-related peptide (CRP-XL) was provided by Richard Farndale (University of Cambridge, UK). Unless stated, all other reagents were from Sigma-Aldrich (Dorset, UK).

### Animals

C57BL/6 mice were purchased from Harlan UK. All animal studies were approved by local research ethics committee (AWERB) and licensed under UK Home Office project license PPL 30/3445.

### Culture and Differentiation of Murine Megakaryocytes

Briefly, bone marrow from C57BL/6 or *Tpm4*^−/−^ mice was isolated and dispersed prior to centrifugation at 200 x g for 10 minutes. Bone marrow was re-suspended in IMDM-Glutamax containing 1% penicillin/streptomycin and 2% serum replacement 1. Cells were cultured for 3 days in the presence of 20 ng/mL rSCF and a further 13 days in the presence of 10 ng/mL rTPO at 37 °C and 5% CO_2_. From day 8 to day 16, cells were transmitted to fresh cell culture dishes every day to reduce megakaryocytes (MKs) to contact with fibroblast cells.

On day 16, MKs were harvested and enriched with a 1.5%-3% albumin gradient for one hour in cell incubator, following stained with either IgG-PE/ or FITC, or CD41-PE or -FITC for 3 hours and further with DNA dye Hoechst 33342 for 20 mins. Then MKs were washed and resuspended in 2 mL medium containing 10 U/mL heparin prior to use.

### Mouse *ex vivo* heart-lung preparation

Mice aged at 8-12 weeks were sacrificed by a Schedule 1 process, by exposure to rising CO_2_. A tracheostomy was then performed, and lungs ventilated with room air or pure nitrogen via a rodent ventilator (Minivent type 845, Harvard Apparatus, USA). Respiratory rate was maintained at 200 breaths/min and tidal volume was 10 mL/kg (~250 μL). The inferior vena cava was then exposed by laparoscopy, and the chest opened by median sternotomy and fat tissue carefully removed.

A 21-gauge catheter was passed into the right ventricle and approx. 100 mL warm (36-38 °C) perfusion buffer (Krebs-Hepes buffer (KHB): 140 mM NaCl, 3.6 mM KCl, 0.5 mM NaH_2_PO_4_,0.2 mM MgSO_4_, 1.5 mM CaCl_2_, 10 mM Hepes (pH 7.4), 2 mM NaHCO_3_) containing 10 U/mL heparin was perfused at 3-4 mL/min into the lung vasculature. Once the colour of mouse lung turned pale, suggesting most of blood in the lung circulation was flushed, the catheter was disconnected.

The left and right superior venae cavae, ascending aorta, inferior vena cava and descending aorta were ligated using 6-0 silk, and a small incision was made in the left ventricle for collection of the perfusate. A suspension of mouse MKs (usually labelled with a fluorescently tagged anti-CD41 Ab) was pumped into the right ventricle, whilst the lungs were either ventilated with air or N_2_ or without ventilation, and the flow rate was maintained at 0.35 mL/min by a SyringeOne Programmable Syringe Pump (Product SKU: NE-1000-ES). The perfusate was collected and re-pumped into the right ventricle after samples were collected for imaging and FACS analysis. The perfusate was recirculated through the pulmonary vasculature in this manner for a total of 18 passages.

After the final passage, 1 mL perfusion buffer was injected into the system to remove some of the remaining cells in the lung vasculature, and all perfusate collected for evaluation of platelet function. The mouse lung was immediately removed and fixed in 4% PFA/PBS. The volume of the lung was measured by fluid displacement.

### Flow cytometry

Platelets derived from IgG-PE stained MKs were set up as negative controls. 25 μL of CD41-PE-stained MK suspension, or perfusates from lung or suspensions from microfluidic chambers after 1, 2, 3, 6, 9, 12, 15 or 18 passages, were fixed in 2% PFA/PBS before FACS. DNA content, viability and mitochondrial membrane potential of generated platelets were determined by Draq5, Calcein AM and TMRM staining, respectively.

To detect the viability and mitochondrial membrane potential of pulmonary endothelial cells (ECs), assays for Calcein Deep Red retention in ECs and TMRM accumulation in active mitochondria were conducted. Pulmonary ECs were isolated from perfused lungs under air-or pure-N2-ventilation or unventilated for approximately 2 hours and marked with either anti-CD31/PECAM-1 or anti-CD102/ICAM-2 antibodies, followed by loading with Calcein Deep Red and TMRM dyes, respectively, for 25 mins at room temperature. Pulmonary ECs from fresh lungs served as control.

Platelet surface glycoproteins were measured by incubating with FITC-conjugated anti-mouse CD61, CD42b, CD42d, CD49b, Glycoprotein VI (GPVI) antibodies and isotype-nonspecific IgG for 20 mins at room temperature before fixation.

To investigate the function of generated platelets (platelets derived from CD41 stained MKs), assays for α_IIb_β_3_ integrin activation (JON/A-binding) and P-selectin exposure were performed. Washed platelets were stimulated with 2 U/ml thrombin or 5 μg/mL CRP for 10 mins followed by incubation with PE-JON/A or PE-P selectin antibodies for 20 mins at room temperature before fixation. Tirofiban and PE-IgG were used as negative controls for the measurement of PE-conjugated JON/A or PE-conjugated P-selectin exposure, respectively.

Samples were analysed on a BD Accuri™ C6 Plus flow cytometer (BD) with 50,000 gated events/sample.

### Two Photon Imaging

After overnight fixation at 4 °C, the lung was staged on a microslicer device (Type: DTK-1000N, UK) and cut into small slices with 800 μm thickness and flat surface. Lung slices from different lobes were fixed into a 100 mm cell culture dish and immersed in PBS for two photon imaging. Imaging was performed using a DeepSee multiphoton laser (Spectra Physics) attached to an upright SP8 confocal microscope (Leica Microsystems). All images were collected using a 25x/0.95NA water dipping lens. For CD41-FITC or CD41-PE, excitation was provided by tuning the multiphoton laser to 927nm, for Hoechst33342, excitation was provided by tuning the multiphoton laser to 750nm. The resultant fluorescence for both scans passing off a SP500 dichroic beam splitter and through both a SP680 filter and either a 525/50 nm bandpass filter to selected only CD41-FITC signal, a 630/75 nm bandpass filter to select only CD41-PE signal or a 460/50 nm bandpass to select only Hoechst33342 signal. CD41-FITC or CD41-PE fluorescence was detected using non-descanned Hybrid detectors (Leica Microsystems) and Hoechst33342 fluorescence was detected using a non-descanned PMT.

Images were acquired with an additional zoom of 2.5x with 1704×1704 pixels (XY) resulting in an effective pixel size of 100 nm. Z stacks were captured with a z-step spacing of 2 μm. All images were capture using a scan speed of 400 Hz with a bidirectional scan.

### Confocal Microscope Imaging

Confocal images were obtained on an inverted SP8 confocal microscope (Leica Microsystems) attached to a DMI6000 microscope frame (Leica Microsystems). All images were acquired using a 100x/1.44 NA oil immersion objective. Excitation was provided by either a 405 nm laser (Hoechst33324) or 488 nm laser (for CD41-FITC) or 561 nm laser (for CD41-PE) and the resultant fluorescence was detected using a Hybrid detector in the range 410-470 nm for Hoechst 33342, 495-570 nm for CD41-FITC or 571-650 nm for CD41-PE. A total of 50 randomly chosen fields of view were imaged and where z-stacks were acquired, a 2 μm z-step spacing was used for Movie S6-8, or a 0.5 μm z-step spacing for Movie S9.

To obtain an estimate of the dimensions of each nucleus in XY, z-stacks were loaded into FIJI and subjected to a maximum intensity projection and a region of interest (ROI) was manually drawn around an individual nucleus. The inbuilt measure function within imageJ was used. To obtain the dimensions of the major and minor axis, the ‘fit ellipse’ option was enabled. To estimate the nuclear depth, analysis was performed in FIJI. Image stacks were loaded, and a cell nucleus was selected using the ROI tool. The average intensity profile of this nucleus was plotted in z and then fitted with a gaussian profile using the built-in plot and fitting tools of FIJI. From the fitted parameters the depth of the nucleus was estimated using the full width at half maximum (FWHM) of the gaussian fit. The FWHM was calculated as 2√(2*ln*2) σ. This process was manually repeated for multiple cell nuclei.

### Microfluidic chamber design, fabrication, and experimental protocol

A set of channels mimicking a physiological vascular system was constructed using standard PDMS approach. The design shows a branching structure such that as branching progressing, the channel diameter decreases by half. All channels are rectangular in cross-section and 10 μm deep, with the largest channels being 100 μm across, decreasing to the smallest channels which are 12.5 μm across. From each larger channel, 16 smaller channels branch off, allowing for maintenance of flow resistance due to the r^4^ power relationship between resistance and channel diameter. Fluid then flows from larger diameter channels to smaller diameter channels, and in reverse on the way out of the system. Cells are passed through the system, repeatedly. The system is scaled up, through multiplexing in parallel, to allow greater cell volumes to be used, as shown in Fig. 1I and Fig. S1C. The mask and SU-8 master mold were fabricated by NuNano (Bristol, UK). The microfluidic channels were fabricated by soft lithography. The mixture of PDMS in a 10:1 ratio was poured over the SU-8 master mold after being degassed in a vacuum desiccator. The PDMS mixture was cured at 80 °C for 2 h and incubated in the oven overnight. The PDMS mold was removed from the SU-8 master. Input (Cell-in) and output (Cell-out) holes were punched using a 0.5 mm OD biopsy puncher (Elveflow, Paris, France). Finally, the PDMS microchannels were irreversibly bonded to glass slides using oxygen plasma for 3 min in a plasma device (Diener Plasma Systems, Ebhausen, Germany).

A suspension of mouse MKs pre-labelled with CD41-PE was pumped into the microfluidic chamber at 0.30 mL/min flow rate. Suspensions were collected from the Cell Out-hole and re-pumped into the microfluidic chamber, after 25 μL samples were taken for FACS analysis. Suspensions were then recirculated through the microfluidic chamber in this manner for a total of 18 passages.

### Transmission Electron Microscope Imaging

To visualize and compare the ultrastructures of generated and control platelets by transmission electron microscopy (TEM), host platelets were first depleted by intraperitoneal administration of anti-GPIbα antibody R300 (2 μg/g bodyweight) prior to MK infusion through the heart-lung preparation. After 18 passages, the generated platelets were pelleted by centrifuging the collected perfusate. The platelet pellet was then resuspended in a cacodylate-buffered glutaraldehyde fixative and fixed at 4-8°C overnight and then post-fixed in osmium ferrocyanide. Fixed cells were then embedded in a solidifying BSA/glutaraldehyde gel. Gel-embedded platelets were stained with uranyl acetate and lead aspartate followed by dehydration with ethanol and infiltrated with Epon resin in a Tissue Processor (Leica EMTP). 70nm sections were cut from blocks with a Reichert Ultracut E and imaged with a Tecnai 12 electron microscope (ThermoFisher UK).

### *In Vitro* Thrombus Formation

In brief, ibidi μ-Slide VI^0.1^ chips were coated with 50 μg/mL collagen overnight at 4 °C before being flushed and blocked with 2% fatty acid–free BSA prepared in HEPES-Tyrode’s buffer. Freshly drawn mouse blood was collected from the inferior vena cava using 10 U/mL heparin and 20 μM PPACK (D-phenylalanyl-prolyl-arginyl chloromethyl ketone) as anticoagulant, following euthanasia by rising CO_2_. Mouse blood was mixed with 2 mL NaCl (150 mM) and centrifuged to remove PRP. Megakaryocytes stained with 2 μM DiOC_6_ were passaged through the mouse-lung preparation 18 times, and approx. 0.7 mL perfusate was mixed with 1.3 mL mouse blood lacking PRP and incubated with CellTracker™ Red CMTPX dye (1:1000 dilution) for 10 mins.

The mixed sample was then transferred to a 5 mL syringe and perfused using an Aladdin AL-1000 syringe pump (World precision instruments, United Kingdom) through the ibidi slide, at a shear rate of 1000/s for 20 minutes. Platelets were fixed by perfusion of 4% paraformaldehyde through channels for 4 minutes before nonadherent cells were flushed away with HEPES Tyrode buffer. Thrombus formation was determined by generating confocal z-stacks (1024×1024 pixels, 0.787 μm z stack distance) from 5 randomly chosen fields of view using a Leica SP8 confocal microscope. Images were acquired using a 20x/0.7 NA air immersion objective. Excitation was provided by either a 488 nm (DiOC_6_) or 561 nm (CellTracker™ Red CMTPX) laser with the resultant fluorescence being detected by Hybrid detectors in the range 498-551 nm (DiOC_6_) or 571-623 nm (CellTracker Red CMTPX).

### Analysis and statistics

All data were analyzed using GraphPad Prism 7 software (GraphPad Software Inc., San Diego, CA, USA). All image analysis was performed using ImageJ-win 64 software. Quantified data are presented as mean ± SEM from at least 3 independent experiments. A value of *P < 0.05 was considered statistically significant and was determined using either a paired Student’s t-test or two-way ANOVA with Dunnett’s or Bonferroni post hoc test for multiple comparison.

---

**Fig. S1.**
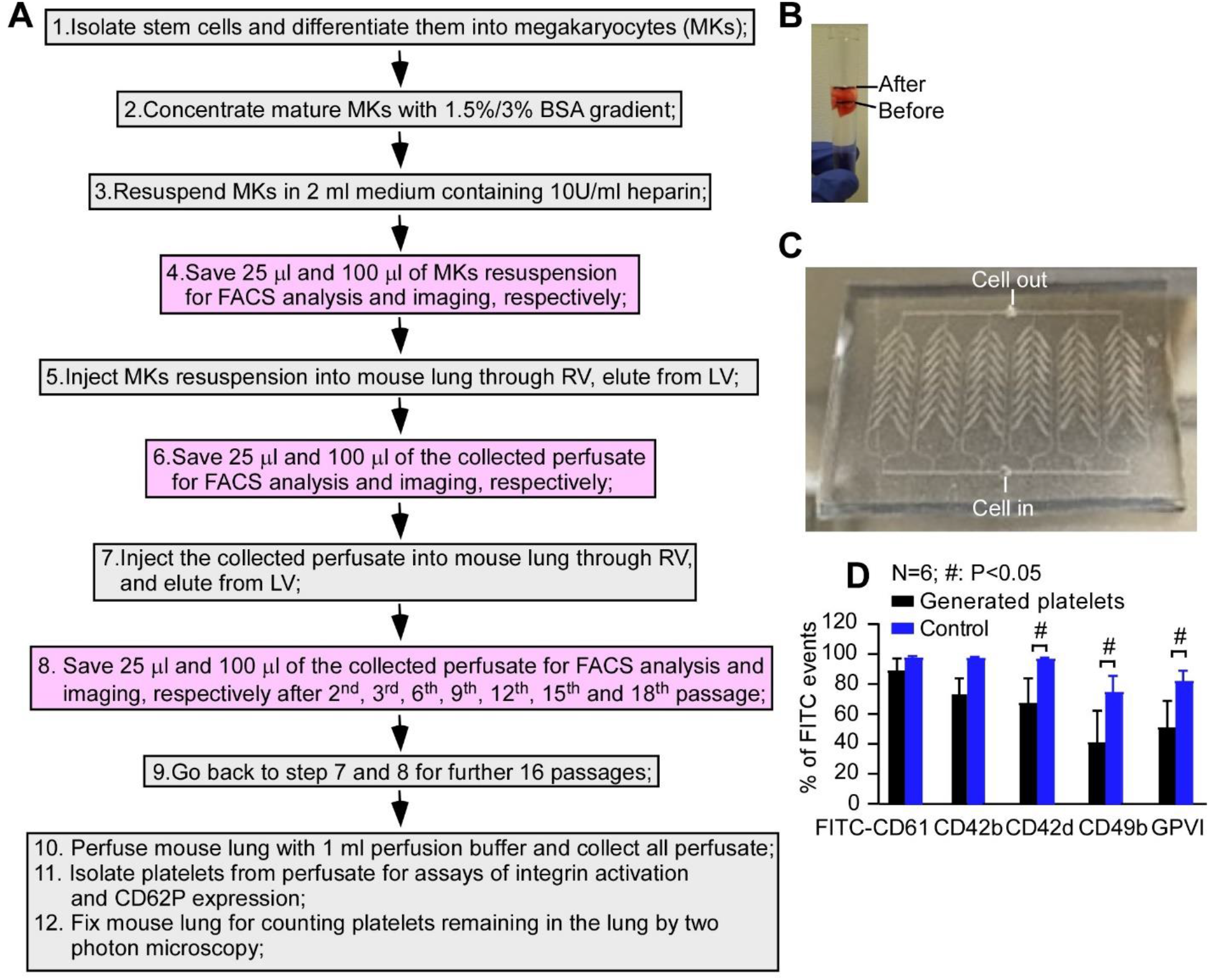
Approach to generating mouse platelets from MKs by infusion through the pulmonary vasculature or a novel microfluidic chamber. **(A)** Experimental protocol for generating platelets from repeated infusion of mouse heart-lung preparation. **(B)** The lung volume after 18 passages was estimated by a fluid displacement technique. This was used to allow accurate estimation of numbers of platelets generated that were still retained within the circulation of the lung. **(C)** Photograph of the microfluidic chamber, made in PDMS by soft lithography, with the arrangement of microchannels visible. Description of the channel architecture is provided in Fig. 1I. **(D)** Platelets generated in the mouse lung vasculature were defined by staining with anti-CD41 PE conjugated antibody. Surface glycoproteins of platelets were measured by FACS on generated platelets and control platelets after staining with different FITC-conjugated antibodies as indicated. Data are presented as % of FITC- positive events. Type or paste caption here. Create a page break and paste in the Figure above the caption.

**Movie S1.**

**3D reconstructed image of *ex vivo* mouse lung – air ventilation.** The demarcation membrane system (DMS) and plasma membrane of mouse MKs were stained with anti-CD41-PE (red) and nuclei stained with Hoechst 33342 (cyan) and passaged through *ex vivo* mouse pulmonary circulation 18 times, whilst lungs were artificially ventilated with air. Images were taken by 2-photon microscopy, and reconstructions made from 20 z-stacks (in 2 μm steps) and reconstructed into 3D space shown in the video. Red events with ~ 2-4 μm in dimension were generated platelets. Images are representative of 4 independent experiments. Scale bar in this video: 20 μm.

**Movie S2.**

**3D reconstructed image of *ex vivo* mouse control lung – air ventilation.** To demonstrate background autofluorescence, lungs were artificially ventilated with air and imaged by 2-photon microscopy, and reconstructions made from 20 z-stacks (in 2 μm steps) and reconstructed into 3D space shown in the video. Images are representative of 4 independent experiments. Scale bar in this video: 20 μm.

**Movie S3.**

**3D reconstructed image of *ex vivo* mouse lung – air ventilation.** Mouse MK DMS and plasma membranes were stained with anti-CD41-FITC (green) and passaged through *ex vivo* mouse pulmonary circulation 18 times, whilst lungs were artificially ventilated with air. Images were taken by 2-photon microscopy, and reconstructions made from 20 z-stacks (in 2 μm steps) and reconstructed into 3D space shown in the video. Images are representative of 6 independent experiments. Green events with ~ 2-4 μm in dimension were generated platelets. Scale bar in this video: 20 μm.

**Movie S4.**

**3D reconstructed image of ex vivo mouse lung – N_2_ ventilation.** Mouse MK DMS and plasma membranes were stained with anti-CD41-FITC (green) and passaged through ex vivo mouse pulmonary circulation 18 times, whilst lungs were artificially ventilated with 100% N_2_. Images were taken by 2-photon microscopy, and reconstructions made from 20 z-stacks (in 2 μm steps) and reconstructed into 3D space shown in the video. Images are representative of 4 independent experiments. Several MKs with very bright green fluorescence are trapped in the pulmonary circulation. Scale bar in this video: 20 μm.

**Movie S5.**

**3D reconstructed image of in vitro thrombus formation under flow.** Generated platelets (stained with DiOC6, plus CellTracker™ Red CMTPX, blue) and control platelets (stained with CellTracker™Red CMTPX alone, magenta) were mixed and perfused through the ibidi slide pre-coated with collagen, at a shear rate of 1000/s for 20 minutes. Images were taken by confocal fluorescence microscopy, and reconstructions made from 30 z-stacks (in 0.787 μm steps) and reconstructed into 3D space shown in the video. Images are representative of 5 independent experiments. Scale bar in this video: 20 μm.

**Movie S6.**

**3D reconstructed confocal image of an intact MK with a central giant lobulated nucleus surrounding by large cytoplasm.** The DMS and plasma membranes of a mouse MK were stained with anti-CD41-PE (red) and nuclei stained with Hoechst 33342 (blue). Images were obtained on an inverted SP8 confocal microscope and 3D reconstruction image was made from 20 z-stacks at 2 μm z-step spacing. Images are representative of 8 independent experiments. Scale bar in the video: 5 μm.

**Movie S7.**

**3D reconstructed confocal image of MK with an extruding giant lobulated nucleus adhering to a large cytoplasmic fragment.** The DMS and plasma membrane of a MK was stained with anti-CD41-PE (red) and nuclei stained with Hoechst 33342 (blue). Images were obtained on an inverted SP8 confocal microscope and 3D reconstruction image was made from 20 z-stacks at 2 μm z-step spacing. Images are representative of 8 independent experiments, Scale bar in the video: 10 μm.

**Movie S8.**

**3D reconstructed confocal image of a naked lobulated nucleus with a completely separated large cytoplasmic fragment.** The DMS and plasma membrane of a MK was stained with anti-CD41-PE (red) and nuclei stained with Hoechst 33342 (blue). Images were obtained on an inverted SP8 confocal microscope and 3D reconstruction image was made from 20 z-stacks at 2 μm z-step spacing. Images are representative of 8 independent experiments, Scale bar in the video: 10 μm.

**Movie S9.**

**3D reconstructed confocal image of sub-nuclei.** The DMS and plasma membrane of MKs were stained with anti-CD41-PE (red) and nuclei stained with Hoechst 33342 (blue) and passaged through *ex vivo* mouse pulmonary circulation 3 times, whilst lungs were artificially ventilated with air. Images were obtained on an inverted SP8 confocal microscope and 3D reconstructed image was made from 20 z-stacks at 0.5 μm z-step spacing. Nuclear lobes were partially fragmenting from a naked MK nucleus. Images are representative of 8 independent experiments. Scale bar in the video: 10 μm.

**Movie S10.**

**3D reconstructed image of *ex vivo* mouse lung – air ventilation.** The DMS and plasma membranes of *Tpm4*^−/−^ MKs were stained with anti-CD41-FITC (green) and passaged through ex vivo mouse pulmonary circulation 18 times, whilst lungs were artificially ventilated with air. Images were taken by 2-photon microscopy, and reconstructions made from 20 z-stacks at 2 μm step spacing and reconstructed into 3D space shown in the video. Abundant fluorescent objects sized 10-12 μm, were visible in focus stacking of 20 continuous 2-photon planes. Images are representative of 3 independent experiments. Scale bar in this video: 20 μm.

